# Coral settlement and recruitment are negatively related to reef fish trait diversity

**DOI:** 10.1101/2021.10.19.464984

**Authors:** Cher F Y Chow, Caitlin Bolton, Nader Boutros, Viviana Brambilla, Luisa Fontoura, Andrew S Hoey, Joshua S Madin, Oscar Pizarro, Damaris Torres-Pulliza, Rachael M Woods, Kyle J A Zawada, Miguel Barbosa, Maria Dornelas

## Abstract

The process of coral recruitment is crucial to the functioning of coral reef ecosystems, as well as recovery of coral assemblages following disturbances. Fishes can be key mediators of this process by removing benthic competitors like algae, but their foraging impacts are capable of being facilitative or harmful to coral recruits depending on species traits. Reef fish assemblages are highly diverse in foraging strategies and the relationship between this diversity with coral settlement and recruitment success remains poorly understood. Here, we investigate how foraging trait diversity of reef fish assemblages covaries with coral settlement and recruitment success across multiple sites at Lizard Island, Great Barrier Reef. Using a multi-model inference approach incorporating six metrics of fish assemblage foraging diversity (foraging rates, trait richness, trait evenness, trait divergence, herbivore abundance, and sessile invertivore abundance), we found that herbivore abundance was positively related to both coral settlement and recruitment success. However, the correlation with herbivore abundance was not as strong in comparison with foraging trait diversity metrics. Coral settlement and recruitment exhibited a negative relationship with foraging trait diversity, especially with trait divergence and richness in settlement. Our findings provide further evidence that fish play a role in making benthic habitats more conducive for coral settlement and recruitment. Because of their ability to shape the reef benthos, the variation of fish biodiversity is likely to contribute to spatially uneven patterns of coral recruitment and reef recovery.

## Introduction

The recovery of coral populations after disturbances, like thermal bleaching and tropical cyclones, depends on larval recruitment, which is known to be heterogeneous across space at local and regional scales (Roff and Mumby 2012; Holbrook et al. 2018; Hughes et al. 2019; Mellin et al. 2019). Studies on coral recruitment outcomes in the field suggest that fish assemblages are an important determinant of recovery trajectories by suppressing key coral competitors like algae through their foraging activities (Korzen et al. 2011; Graham et al. 2015; Kuempel and Altieri 2017).

Dynamics at the early life stages of coral settlement and recruitment are critical bottlenecks in the recovery of reef coral assemblages from disturbances (Ritson-Williams et al. 2009; Adjeroud et al. 2017). Settlement refers to the life stage where planktonic coral larvae establish onto substrates as sessile spat. Recruitment occurs when spat form coral colonies through growth. Both of these life stages are marked by high mortality rates (Vermeij and Sandin 2008). Successful coral settlement requires optimum water flow conditions as well as available substrate space (Chadwick and Morrow 2011; Hata et al. 2017). On the other hand, recruitment success involves competing with other benthic organisms for resources and light as well as avoiding predation (Doropoulos et al. 2016). The ability to survive and compete for space are strong determinants of survival for corals in these early life stages.

Algae are major competitors with corals for space and resources. Their specific competitive mechanisms differ according to morphological groups. Upright foliose macroalgae outcompete corals primarily through shading effects (Webster et al. 2015), while lower profile morphologies like turfing and encrusting algae compete through space pre-emption and maintaining unfavourable sedimentation conditions (Wakwella et al. 2020). Algae are able to proliferate quickly in response to space availability, as demonstrated by rapid colonisation of algae following massive coral community mortalities (McCook et al. 2001; Kuffner et al. 2006; Diaz-Pulido et al. 2010). Because of their fast growth, algae can often quickly dominate coral reefs and inhibit coral replenishment (Hughes 1994; McClanahan et al. 2001; Rogers and Miller 2006; Bruno et al. 2009; Clements et al. 2018; Bozec et al. 2019). The ability of coral reef ecosystems to balance algal productivity without overgrowth has largely been attributed to foraging by herbivorous reef fishes (Graham et al. 2013; Kuempel and Altieri 2017; Manikandan et al. 2017; Dajka et al. 2019), which collectively have been estimated to consume up to 65% of net primary productivity on a reef (Polunin and Klumpp 1992). By suppressing the standing biomass of algae, herbivorous fishes are often considered indirect facilitators of coral settlement and recruitment (Bellwood et al. 2006; Hughes et al. 2007; Chong-Seng et al. 2014; Doropoulos et al. 2017).

The foraging impact from fishes on the benthic assemblage is mediated by their behavioural and physical characteristics (functional traits). Not all bites are equal in the removal of algal biomass, and some can even be destructive to corals, both recruits and adults (Baria et al. 2010; Evans et al. 2013; Bonaldo and Rotjan 2017). Trait-driven variation in foraging impacts can be assessed at two scales: among species and among assemblages. Foraging impacts among species vary according to traits such as food selectivity, jaw morphology, and biting mode, which are often summarised in functional groupings especially for herbivorous fishes (Mantyka and Bellwood 2007; Green and Bellwood 2009; Michael et al. 2013; Streit et al. 2015, 2019). Food selectivity is especially relevant as fish species target algae differentially, from sediment load reduction in detritivores (Goatley and Bellwood 2010; Tebbett et al. 2017), macroalgae removal in browsers (Hoey and Bellwood 2009; Tebbett et al. 2020) to total removal of turf by croppers and scrapers (Korzen et al. 2011).

Specifically in the context of early life-stage survival in corals, trait-based analyses have pointed to important species-driven differences in foraging. Parrotfishes, with their beak-like dentition, have scraping and excavating foraging modes, and as such they can induce coral recruit mortality through intense benthic interactions (Penin et al. 2011a; Bonaldo and Rotjan 2017). There is also considerable variation within functional groupings. For example, most rabbitfishes are typically categorised as ‘algal croppers’ yet there is evidence that several species (e.g., *S. puellus, S. punctatus, S. punctatissimus)* have diverse diets that include benthic invertebrates (Hoey et al. 2013). Studies have also shown that topographic refuges play a critical role in recruitment success as they physically prevent more disruptive foragers from interfering with the coral recruitment process (Doropoulos et al. 2012; Brandl and Bellwood 2016; Gallagher and Doropoulos 2017). Hence, the balance between positive and negative foraging impacts on coral recruitment from fish assemblages depends on the trait composition as well as their respective benthic environments.

Other benthic taxa (e.g. sponges) also compete with corals and point to the need to consider the effects of other benthic foragers on coral settlement and survival (Elliott et al. 2016; Madduppa et al. 2017). For example, sessile invertivores may also lend a similar facilitative effect to corals by suppressing other benthic competitors, such as sponges and soft corals. It is not yet clear what, if any, effect invertivores have on coral settlement and recruitment.

Foraging impacts, whether beneficial for space-clearing or harmful to corals, vary with species traits, therefore impacts delivered collectively by a fish assemblage would vary according to the distribution and composition of these traits (Cheal et al. 2010). The species and trait composition of fish assemblages vary widely across space in coral reefs, depending on structural complexity of the habitat and environmental gradients (Cheal et al. 2012; Darling et al. 2017; Richardson et al. 2017; Bach et al. 2019). Trait variation within an assemblage results in highly differentiated strategies between species (trait complementarity) and similar overlapping strategies (trait redundancy). Foraging trait complementarity between specialist species has been shown to be most effective at reducing algal cover for coral juveniles (Burkepile and Hay 2008, 2011). However, this pattern may not be general as a considerable number of herbivory studies have also shown that key species uphold a majority of this function (Bellwood et al. 2006; Hoey and Bellwood 2009; Vergés et al. 2012; Michael et al. 2013; Tebbett et al. 2020). These studies suggest that a small number of species may be disproportionately influencing reef benthos irrespective of the fish assemblage diversity, which is a pattern also detected in other consumption functions across tropical reefs (Schiettekatte et al. 2022). It is also unclear how variation in fish assemblage foraging-relevant traits links with spatial patterns in coral recruitment. Furthermore, trait diversity effects in foraging impacts have not yet been investigated beyond assessing effects related to functional groupings as a proxy of traits (Brandl et al. 2019).

Here, we investigate whether variation in foraging trait diversity of fish assemblages correlates with variation in coral settlement and subsequent recruitment to juvenile cohorts. Given previous evidence of positive species diversity effects on herbivory (Burkepile and Hay 2008; Rasher et al. 2013) and the positive scaling of trait richness with species richness, we hypothesise that coral settlement and recruitment will be more successful where there are more trait diverse fish assemblages. Specifically, we examine whether greater foraging rates, trait richness, trait evenness, trait divergence, herbivore abundance, and benthic invertivore abundance are associated with coral settlement and recruitment success.

## Materials and methods

### Study location

We conducted the study at seven sites (1.4–3.7 m depth) representative of the variation in topography and abiotic substrate within a no-take marine national park zone at Lizard Island (14°40’ S, 145°28’ E) in the northern Great Barrier Reef, Australia (Figure 1). Recent coral mortalities from thermal bleaching and cyclone damage observed at Lizard Island (Madin et al. 2018; Hughes et al. 2019) made this an opportune time and location to investigate coral settlement and recruitment dynamics post-disturbance. Data collection took place during early austral summer surrounding the annual spawning event, from November-January. Coral data were collected in 2018-19 and 2019-20, and fish assemblage data were collected in 2019-20.

**Figure 1.**
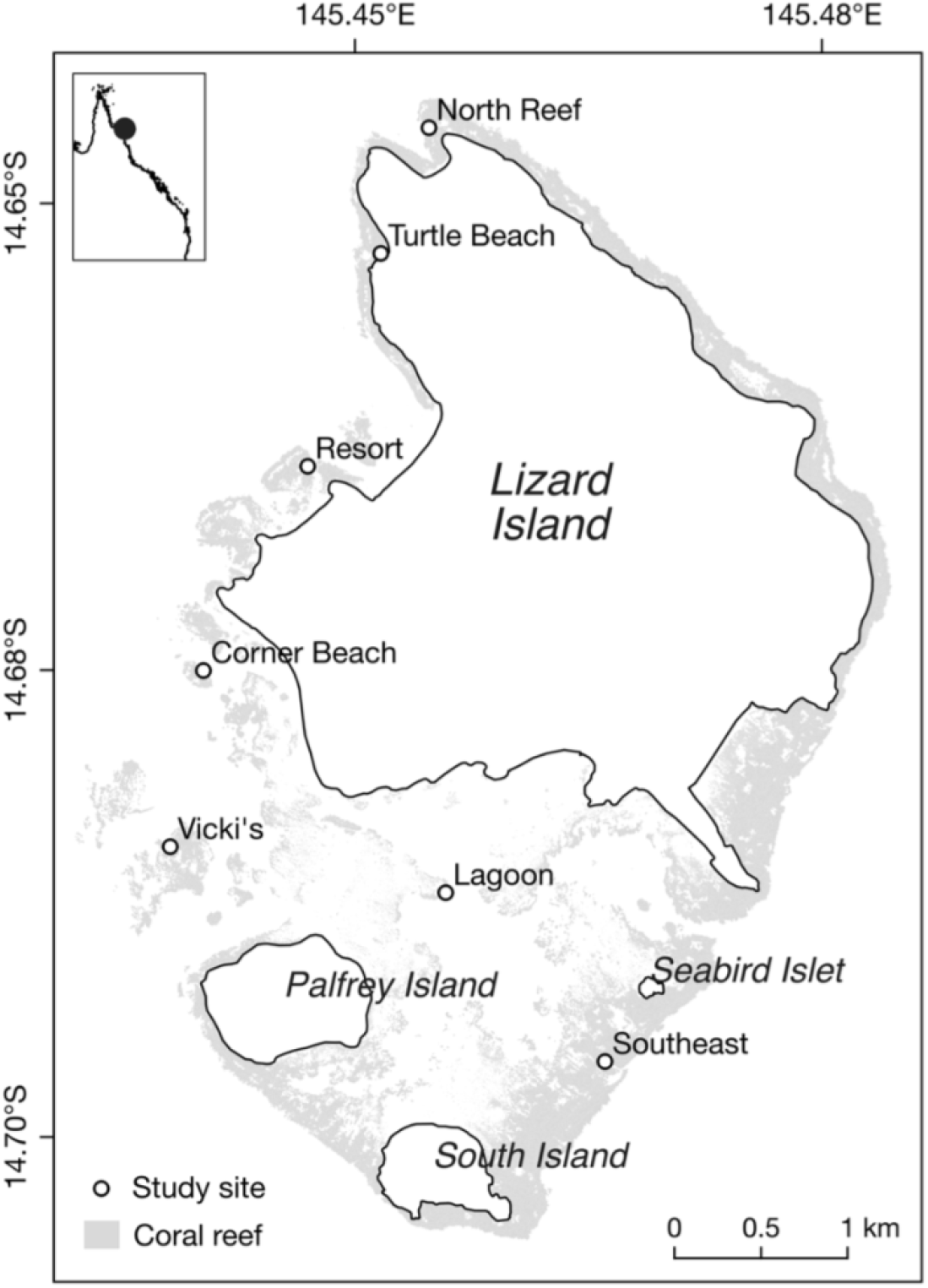
Map of study site locations around Lizard Island. The coral reef area is shown shaded in light grey. Spatial data for reef and coastline boundaries were sourced from the Great Barrier Reef Marine Park Authority Geoportal (GBRMPA 2020) and Roelfsema et al. (2014).

### Remote underwater videos

We obtained fish assemblage and bite data from remote underwater videos (RUVs), using an adaptation of baited remote underwater video methods (Langlois et al. 2018). At each site, we deployed a single waterproof camera (GoPro Hero4 Session on a wide setting) in acrylic housing on an abiotic substrate. We placed markers at a 2 m radius from the camera lens, establishing a sampling area of approximately 4 sq m with a camera field of view measuring 118°. Deployment lasted for a total of 45 minutes at each site, with the first 15 minutes omitted from processing to avoid diver and boat presence influencing observations.

We processed videos in two iterations, the first to count and identify individual fish within the marked sampling area to species level or lowest possible taxon when possible, and the second to enumerate foraging rates. Observation records where we could not identify to the genus level with certainty were omitted from analysis. To reduce potential bias of double counting highly site attached fish, we identified recurring individuals of the same species and relative size that had been previously observed in the same location with similar behaviours. For bite data, we recorded total bites and length class of the individual fish biting. We used visual estimation for fish length classifications (< 5 cm, 5–9 cm, 10–19 cm, 20-29 cm, and so on in 10 cm intervals inclusive). We then recorded total in-frame occurrence time for all species at the site observed biting at least once, regardless of the behaviour during the occurrence. Unlike processing for fish assemblage structure, this bite observation did not distinguish between individuals.

### Coral settlement and recruitment

To quantify coral settlement, we sampled settling coral spat using experimental substrates in the summers of 2018-19 and 2019-20. In both years, six unglazed clay tiles (11 × 11 cm) were deployed horizontally onto permanent mountings installed at each site (*n* = 42). We deployed tiles one week before predicted coral spawning to allow for establishment of biofilms and crustose coralline algae that reflect the natural conditions of available hard substrate on a reef (Heyward and Negri 1999). We collected tiles after two months and subsequently bleached and dried them for inspection under dissection microscope to count coral spat.

We counted coral recruits *in situ* aided by georeferenced orthomosaic reconstructions of 100 m^2^ reef areas (“reef records”) at each site. Recruits were defined as new colonies which were not fragments of previous colonies and had ≤ 5 cm in diameter (Bak and Engel 1979). These orthomosaics were produced from photogrammetric models following the pipeline of Pizarro et al. (2017) as adapted by Torres-Pulliza et al. (2020). We divided orthomosaics into quadrants for each site (*n* = 28), which were then annotated *in situ* with location and identification for all recruit and adult coral colonies. We identified recruits in 2019 by comparing annotation changes from 2018.

### Fish assemblage predictor variables

We compiled six foraging traits for the fish species observed in RUVs. These traits were selected to represent assemblage diversity with respect to foraging ecology, interactions with substrate, substrate impact, and foraging range (Table 1). Using trophic and diet data from FishBase extracted with the rfishbase R package (version 3.0.4; Boettiger et al., 2012) we assigned trophic groupings according to the definitions established by Parravicini et al. (2020). We also used diet and food item data to allocate the water column position of foraging (benthic, demersal, pelagic/mid-water). If a majority of food items within the diet were specified to be benthic substrata or zoobenthos, we assigned a category of benthic foraging. Where diets consisted of a minority of food items found on the benthos, we classified as demersal. Exclusive planktivores and piscivores we assigned as mid-water/pelagic foragers. Foraging mode groupings were based on the classifications outlined by Green and Bellwood (2009), Cheal et al. (2010), and Stuart-Smith et al. (2013). Details on assigning foraging mode categories are described in the supplementary material.

**Table 1.** Traits used to quantify the functional diversity of reef fish assemblages in regard to feeding ecology, substrate interaction, and delivery of feeding functions. Values were extracted or derived from various databases and literature.

| Trait | Type | Levels/units | Source |
| --- | --- | --- | --- |
| Functional group | Factor | Herbivore/microvore, detritivore, planktivore, corallivore, microinvertivore, macroinvertivore, crustacevore, sessile invertivore, piscivore | 1, 2, 3 |
| Foraging mode | Factor | Excavator, cropper, scraper, browser, brusher, picker, farmer, suction feeder, ambush feeder, active feeder | 1, 4, 5 |
| Trophic level | Continuous | 2.0–5.0 | 1 |
| Water column position of feeding | Factor | Pelagic, demersal, benthic | 1 |
| Residency/Range | Ordered factor | Index of residency and active range, 1-5 with 1 representing highly territorial species and 5 for wide-ranging pelagic species | 1, 7-11 |
| Schooling | Ordered factor | Index of schooling behaviours during feeding from 1-4, with 1 representing solitary species to 4 being species forming large shoals or schools | 1, 6 |
| 1. Pauly and Froese 2019 |  | 5. Purcell and Bellwood 1993 | 9. Welsh and Bellwood 2012 |
| 2. Parravicini et al. 2020 |  | 6. Randall et al. 1996 | 10. Pillans et al. 2014 |
| 3. Brandl and Bellwood 2014 |  | 7. Meyer and Holland 2005 | 11. Davis et al. 2015 |
| 4. Green and Bellwood 2009 |  | 8. Meyer et al. 2010 |  |

Following classification of fish species functional groupings, we calculated the relative abundance of herbivores/microvores (Clements et al. 2017) and sessile invertivores for assemblages at each site.

### Trait diversity analysis

To assess and compare the foraging trait diversity of fish assemblages, we generated three complementary indices of 1) trait richness via the trait onion peeling index (TOP; Fontana et al. 2016), 2) trait evenness, and 3) trait divergence (see Villéger et al. 2008) from a global trait space. TOP quantifies the volume of the trait space filled by the assemblage, where higher measures indicate that the assemblage occupies more trait space and hence richer in traits. TOP is the sum of convex hull volumes calculated by sequentially eliminating species at vertices, hence “onion peels” of convex hulls (Fontana et al. 2016; Legras et al. 2018). Trait evenness describes the variation in distance in the trait space between adjacent species, where higher measures of evenness mean that the abundance of species within an assemblage are more equally distributed throughout the trait space. Lastly, trait divergence measures the distribution of an assemblage relative to the trait space centroid and extremes. Higher trait divergence values reflect greater trait differentiation between species and therefore indicates an assemblage with very little trait overlaps or redundancy. Both evenness and divergence are weighted by species abundance.

Construction of the trait space was performed using Principal Coordinates Analysis (PCoA) based on Gower dissimilarities between all species observed in our study according to the six foraging traits (Villéger et al. 2008; Laliberté and Legendre 2010). Ordered factor traits were handled using the Podani method (Podani 1999) and Cailliez corrections to conform the matrix to Euclidean space, which prevents the generation of negative eigenvalues during scaling (Legendre and Legendre 2012). The resulting trait indices are orthogonal, and so correlation between any of these measures are not due to mathematical artefacts but rather to characteristics of the assemblages (Mason et al. 2005). Dissimilarity, trait space construction, trait evenness, and trait divergence calculations were all performed with the FD R package (Laliberté et al. 2014). TOP was calculated using code provided in Fontana et al. (2016).We also quantified the relative contribution from individual species to the trait diversity of each site using a “leave one out” approach. For each species, we omitted its dissimilarities from the dissimilarity matrix, then used this matrix to reconstruct the trait space and recalculate trait diversity indices. We calculated the species contribution to each site’s trait indices as:

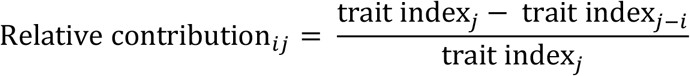

Which is the difference between the original index measure *j* from the omission index measure *j – i* divided by the original index measure. Hence, a positive relative contribution means that the inclusion of a species resulted in a greater trait index and vice versa.

### Calculation of site-level foraging rates

We standardised bite counts by the total observation time for each species to give bite rates (bites min^-1^) for each length class at each site. As our goal was to calculate a foraging rate at the site-level from total bites observed, we did not standardize by number of biting fish. Bite rates were then aggregated by length class. To factor the difference in foraging impacts (i.e. substrate removal) due to trophic group, foraging mode, and water column position traits (Purcell and Bellwood 1993; Green and Bellwood 2009; Burkepile and Hay 2010; Hoey and Bellwood 2011), we calculated a species trait-based coefficient to scale bite rates (details in Supplementary Material). To factor the difference in foraging impacts due to differences in fish size (and hence bite sizes, see Adam et al. 2018; Hoey 2018), we scaled bite rates by the size class midpoint length for individuals of each length class (e.g. 7.5 cm for length class 5–10 cm). We then obtained a foraging rate (bites-cm min^-1^) for each site following Equation 1, where S*_i_* is the trait-based coefficient for species *i*, *L_il_* is the median length for individuals in length class *l* of species *i*, and B*_il_* the bite rate by length class and species for each study site. We refer to bite rates as foraging rates (in bites-cm min^-1^) after this scaling. Given the utility of this foraging rate for relative comparison and not for an objective quantity, we then scaled foraging rates by their standard deviation to place it on a common effect size scale with other explanatory variables for ease of interpretation, as they were indices or proportions constrained between 0 and 1.

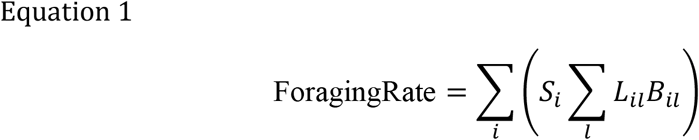

### Statistical modelling and sensitivity analyses

We modelled coral settlement and recruitment through spat and recruit counts respectively as functions of six predictors that captured realised and potential foraging impact. Only 2019-20 coral data were used as response variables in our modelling. Foraging rates represented realised foraging impacts while trait richness (TOP), evenness (TEve), divergence (TDiv), herbivore abundance (Herb), and sessile invertivore abundance (SessInv) represented potential foraging impacts. Site was included as a random intercept term to account for non-independence in same-site coral abundances (Equations 2 and 3).

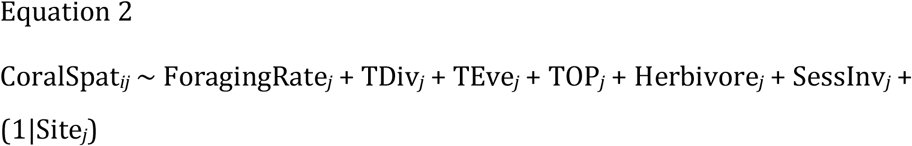

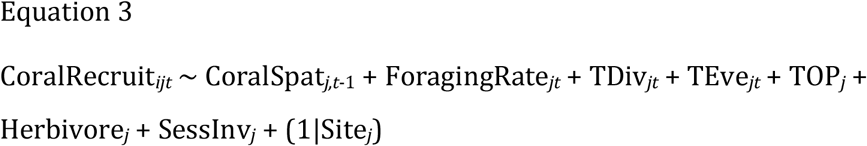

Predictor variables for the coral settlement model are expressed for each site *j* while the coral spat counts exist per settlement tile *i* grouped in six per site *j* (Equation 2). The recruitment model is structured similarly where *i* represents a recruitment quadrant at site *j* (Equation 3). For recruitment specifically, we also included coral spat counts from 2018-19 (*t*–1) as an explanatory covariate to account for the way recruitment could be limited by settlement rates the year prior. All other predictor variables for time *t* refer to 2019-20. We checked for collinearity between predictor variables using Pearson correlation coefficients prior to model fitting. Due to predictor variables reflecting various aspects of a shared fish assemblage at each site, we accepted correlation coefficients between predictors below 0.8 (Figure S1).

To determine the most parsimonious effect structure that captures settlement and recruitment patterns, we used a multi-model inference approach for the response variables of coral spats and recruits. We fitted a full generalised linear mixed model with negative binomial errors and log link function for each response variable using the lme4 package (version 1.1-23, Bates et al., 2015). All above analyses were conducted in R 4.0.0 (R Core Team 2020).

From the full models described above, we constructed two sets of candidate models with all possible combinations of potential foraging impact fixed effects. All candidate combinations included foraging rates. Our null model consisted of no fixed effects, only site as a random intercept term. We also fitted a second null model for coral recruitment consisting of 2018 spat counts as a fixed effect and again, site as a random intercept. We ranked all candidates using Akaike Information Criterion values corrected for small sample sizes (AICc) for model selection (Burnham and Anderson 2002). Selection of the optimum coral spat and recruit models was based on the lowest AICc value (MuMIn package; Bartoń 2020). We also calculated AICc weights as estimates of the probability that each model is the optimum candidate. If top-ranked models were within a difference of 2 AICc, we selected the candidate with a greater AICc weighting. If AICc weights could not differentiate model candidates, we then used residual deviance as a tie-breaker.

We conducted two sensitivity analyses to assess whether our sampling effort was consistent in capturing the local fish assemblage composition. To assess whether the duration of our sampling effort was sufficient, we calculated cumulative species counts for every timestamp where we observed fish individuals. For each site, we then fitted asymptotic and Gompertz regression models to the species accumulation curves to examine whether saturation was achieved within 30 minutes. To assess our sampling area, we compared fish assemblage data for a subset of our sites with additional observations from secondary backup video footage, specifically in trait space construction both independently and combined. We deployed two backup video cameras at all sites in the event of recording failure or changes in camera positioning due to wave exposure or fish activity. Each camera captured a different sampling area in the study site. We had viable video footage from one backup camera matching the sampling duration for three sites (North Reef, Turtle Beach, and Southeast). We were able to select video segments for North Reef and Turtle Beach temporally separate from the original videos to minimise the influence from highly mobile individuals appearing in multiple cameras at similar times. We first checked if PCoA results independent of the original data returned similar scaling for trait space. We then visually compared the overlap of assemblages within a common trait space and calculated Bray-Curtis dissimilarity to quantify assemblage composition differences (vegan R package, Dixon 2003).

## Results

We identified a total of 624 individual fish from 104 species from a total 3.5 hours of video recordings. Fish abundance across the seven study sites ranged from 37 individuals at Turtle Beach to 210 at Southeast, with an overall mean of 89 ± 66 individuals SD. The 104 species observed were dominated by herbivores (33.7%) and macroinvertivores (14.4%). Overall, the relative abundance of herbivores was 32.5% ± 17.6% SD and ranged from 8.1% in Turtle Beach to 62.4% in Southeast (Figure 2). The mean relative abundance of sessile invertivores was lower in contrast at 1.3% ± 1.5% SD (Figure 2).

**Figure 2.**
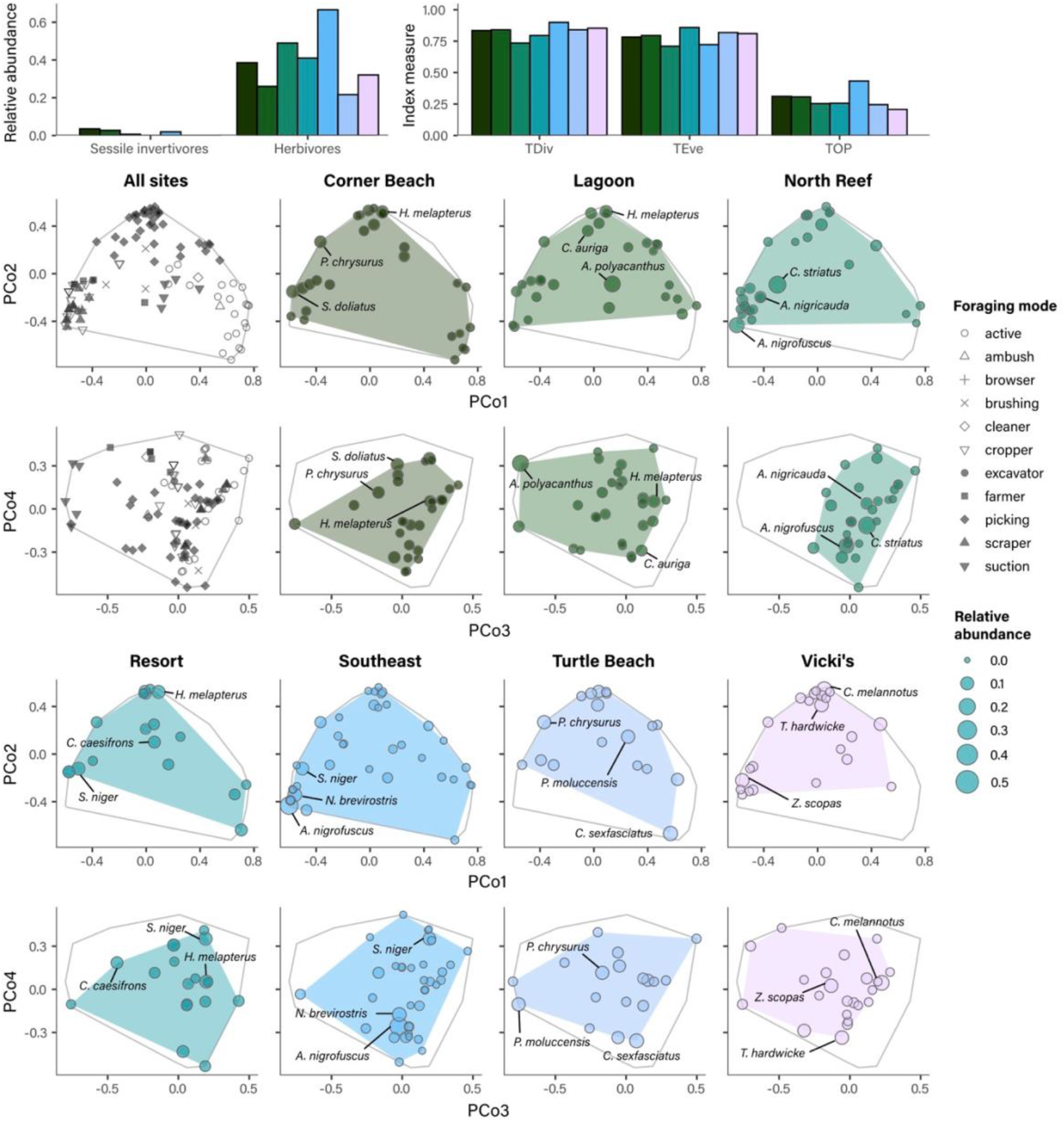
Trait diversity of fish assemblages at the site-level. The bar graphs (top) show measures for relative sessile invertivore and herbivore abundance (top left) and trait diversity indices (top right): trait divergence (TDiv), trait evenness (TEve), and trait onion peeling index for trait richness (TOP). These three facets of trait diversity relate to the volume of the occupied trait space (TOP; i.e. trait richness), the regularity of species distributed within the space (TEve), and the dispersion of the assemblage towards the trait extremes of the space (TDiv). The array shown is a four-dimensional representation of assemblages according to six foraging traits of species. Species are represented by circles, with varying sizes by relative abundance. Distance between circles represents trait dissimilarity between species. The trait space occupied by the assemblage is shaded to represent TOP. For comparison, the reef-level trait space (i.e. all sites, representing TOP = 1) is shown as a grey outline.

### Trait space and trait diversity metrics

The resulting four-dimensional global trait space captured 36.6% of the variation (i.e. proportional sum of eigenvalues; Figure S2a). Our validation of preserved trait space dissimilarities in the Mantel test returned a significant strong correlation (rM = 0.868, *p* < 0.01; Figure S2b). Detritivores and planktivores were located toward the centre of the trait space in the first two dimensions (Figure S6 in Supplementary Material) while herbivores clustered tightly in the lower left corner and corallivores in the upper middle corner. In contrast, large differences in trait richness in the third and fourth dimensions were driven by solitary species with small active ranges and schooling species with large active ranges (Figure S7). Trait richness was relatively similar across sites apart from a notable outlier in Southeast (TOP = 0.43), ranging from 0.20 at Vicki’s to 0.31 at Corner Beach (Figure 2). The fish assemblage composition at Southeast contained relatively more trait extreme species in all four dimensions (Figure 2, Table S2), resulting in the lowest trait evenness measures (TEve = 0.72) and greatest trait divergence (TDiv = 0.90). In contrast, the assemblage at North Reef was abundant in centrally clustered species and hence the least trait divergent (TDiv = 0.74; Figure 2).

### Sensitivity analyses

Species accumulation curves showed that while sites differed in accumulation rates (i.e. time of saturation), all sites were sufficiently saturated at the end of the 30-minute sampling duration (Figure S3 in Supplementary Material). Trait space comparisons with our backup assemblage data demonstrated a high degree of overlap and there were no significant additions to the assemblage when these sample areas were pooled (Figure S4a). Bray-Curtis dissimilarity indices for the sites of North Reef, Southeast, and Turtle Beach ranged from 0.360 to 0.688 and mean change in trait diversity indices was 0.039 ± 0.068 SD. While there were larger differences in TOP driven by some trait-extreme species, especially in North Reef, the relative rankings between sites were preserved (Figure S4b). Trait space construction of the two sets of videos did not show significantly different mappings of species within the assemblage and site-wise differences in trait diversity metrics remained remarkably consistent (Figure S5). Given the evidence from these analyses, we find that our sampling effort both in space and time were sufficient to capture fish assemblage diversity at Lizard Island.

### Foraging rates

35 fish species were observed biting the substrata. Resulting trait-weighted coefficients to reflect bite impact ranged from 0.05 for suction-feeding planktivores to 3.67 for excavator herbivores (Table S1). Five dominant biting species contributed to more than 50% of the total foraging rates observed at sites: *Ctenochaetus striatus* (15.4%), *Chlorurus spilurus* (12.6%), *Hemigymnus melapterus* (8.9%), *Chlorurus microrhinos* (8.6%), and *Acanthurus nigrofuscus* (7.6%). Herbivores, mainly excavators and algal croppers, were the most intense foragers especially at the sites Corner Beach, North Reef, and Vicki’s, even though they were not the most prevalent (Figure 3).

**Figure 3.**
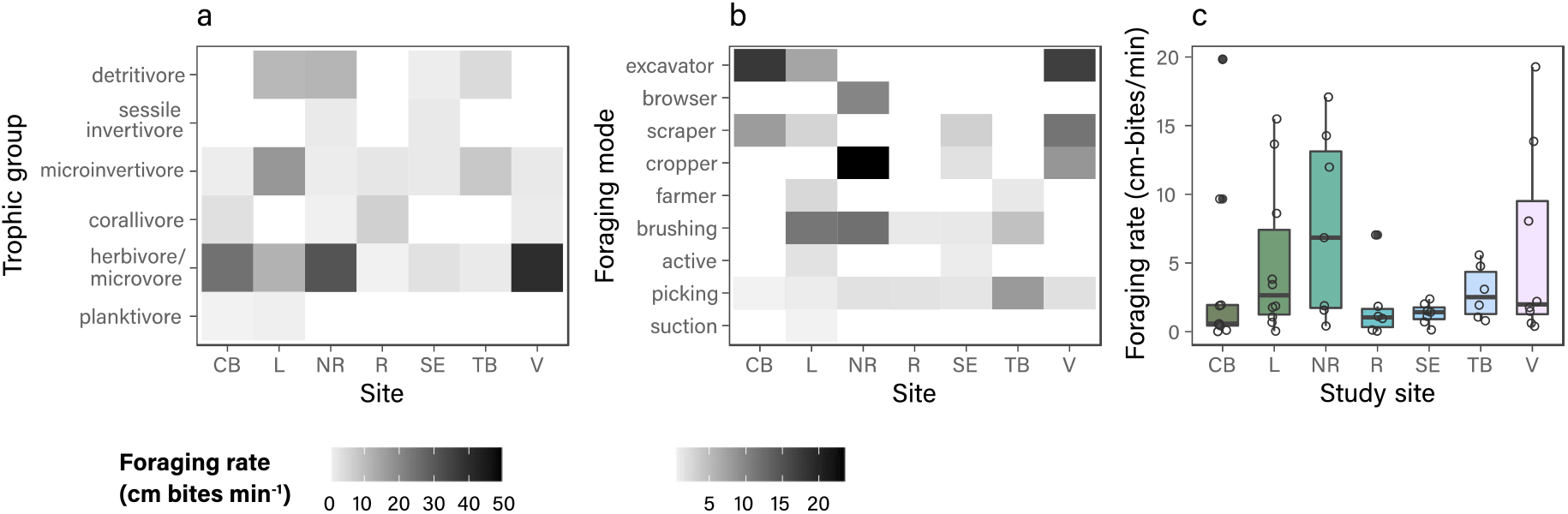
Observed foraging rates at each study site, Corner Beach (CB), Lagoon (L), North Reef (NR), Resort (R), Southeast (SE), Turtle Beach (TB), Vicki’s (V). Foraging rates (cm bites min^-1^) are grouped according to contributions by trophic group (a) and foraging mode (b). Both panels (a) and (b) represent foraging rates by shading, where darker shading represents higher feeding rates and vice versa. Note differences in scales as foraging rates range from 0.03-43.4 (a) and 0.03–23.9 (b). White represents absent groups from sites. Overall foraging rate distributions for species in each site are shown in (c).

### Coral settlement and recruitment

Coral settlement and recruitment reflected similar patterns across our study sites (Figure S8). Settlement was consistently low at Lagoon, Southeast, and Corner Beach (Figures 4-5), ranging from 3-18 total spats summed across six tiles in 2018-19 and 8-14 spats in 2019-20. Coral recruitment was low at Lagoon (mean of 4.00 colonies ± 4.00 SD) and Turtle Beach (8.25 colonies ± 12.53 SD; Figures 4-5; Figure S8). Both coral settlement and recruitment in 2019-20 were highest at North Reef, where there was an average of 13.83 spats per settlement tile ± 6.52 SD (total of 83 spats) and 57.25 recruit colonies per site quadrant ± 21.69 SD (Figures 4-5; Figure S8).

**Figure 4.**
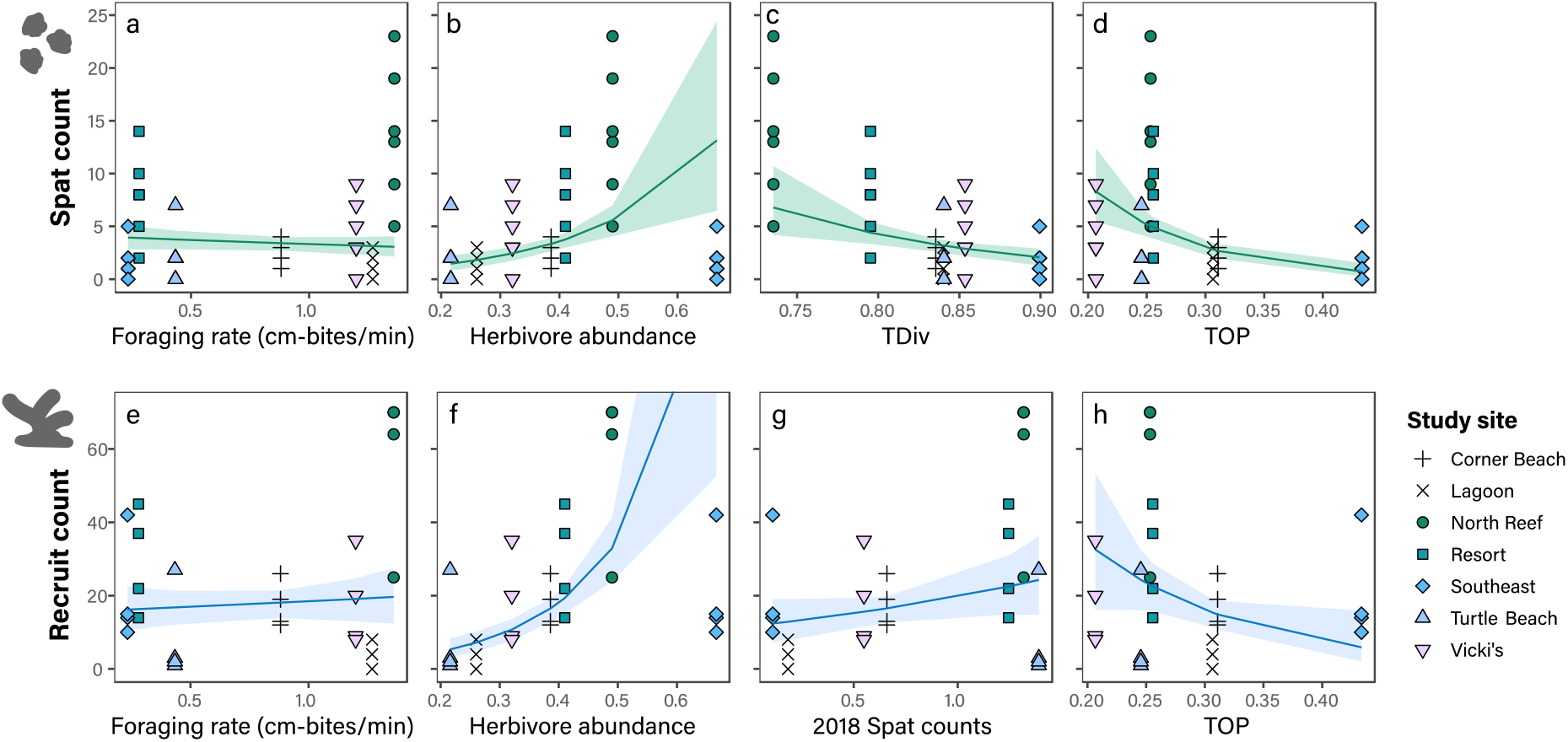
Partial predictions for the optimum models relating coral settlement (above) and recruitment (below) with fish assemblages. The settlement model includes the fixed effects of (a) scaled foraging rate of fishes (cm bites min^-1^), (b) relative herbivore abundance, (c) trait divergence, and (d) trait richness (TOP). The recruitment model includes the fixed effects of (e) spat abundances from 2018, (f) scaled foraging rate of fishes (cm bites min^-1^), (g) relative abundance of herbivorous fish, and (h) trait richness (TOP). Both spat counts and foraging rates were scaled by their range. Coral spat were counted from six settlement tiles at seven sites *(n* = 42) and coral recruits were counted from four quadrants of each circular study site (*n* = 28). Each data point here represents a tile or a quadrant grouped by site in various shapes. Partial predictions from the model for each parameter are represented by solid coloured lines with bootstrapped confidence intervals (from 999 simulations) shown shaded.

### Optimum predictors of coral settlement and recruitment

The fixed effect structure that best explained variation in coral settlement consisted of foraging rate, trait divergence, TOP, and herbivore abundance (Table 2). Although we identified strong negative correlation between herbivore abundances, TOP (*r* = 0.69) and trait evenness (*r* = −0.67) in pairwise checks, most model candidates including trait evenness did not perform well (Table 2). Interestingly, coral settlement and recruitment only differed in trait divergence in their optimum fixed effect structures. Coral settlement was best explained by foraging rates, trait divergence, trait richness, and herbivore abundance (Table 2). This top-ranking settlement model candidate performed markedly better than other candidates (ΔAICc = 2.99, Table 2), but model selection was not as clearly distinguished between recruitment model candidates. Three highest ranking recruitment model candidates fell within less than 0.25 ΔAICc, all including herbivore abundance but varied in the inclusion of trait diversity predictors (Table 2). From our tiered ranking criteria, the final selected recruitment model included 2018 spat counts, foraging rates, herbivore abundance, and trait richness (Table 2).

**Table 2.** Ranking of top candidate and null models for coral spat and recruit models. Site is included in every candidate as a random intercept term, represented as (1|Site). Fixed effect structures vary in fish assemblage diversity variables of trait divergence (TDiv), trait evenness (TEve), trait richness (TOP), herbivore abundance (Herb), and sessile invertivore abundance (SessInv). All candidates include foraging rates (For) and, for recruit models, spat counts from 2018. Model candidates were ranked according to their AICc values. Top-ranked models are bolded for emphasis. Candidates that failed to converge were omitted.

| Models | AICc | $\Delta\text{AICc}$ | Weight | mR<sup>2</sup> | dev |
| --- | --- | --- | --- | --- | --- |
| *Coral settlement* |  |  |  |  |  |
| **For + TDiv + TOP + Herb + (1\|Site)** | **202.74** | **0** | **0.488** | **0.639** | **185.44** |
| For + TDiv + TEve + TOP + Herb + (1\|Site) | 205.73 | 2.99 | 0.110 | 0.638 | 185.37 |
| For + TDiv + TOP + Herb + SessInv + (1\|Site) | 205.80 | 3.06 | 0.106 | 0.638 | 185.44 |
| For + TOP + Herb + (1\|Site) | 206.50 | 3.76 | 0.075 | 0.634 | 192.10 |
| For + TDiv + SessInv + Herb + (1\|Site) | 206.53 | 3.79 | 0.073 | 0.566 | 189.24 |
| For + TDiv + TEve + Herb + SessInv + (1\|Site) | 207.07 | 4.33 | 0.056 | 0.638 | 186.70 |
| (1\|Site) | 213.20 | 10.46 | 0.003 | 0 | 206.56 |
| *Coral recruitment* |  |  |  |  |  |
| **Spat2018 + For + Herb + TOP + (1\|Site)** | **222.62** | **0** | **0.23** | **0.586** | **202.73** |
| Spat2018 + For + Herb + (1\|Site) | 222.63 | 0.01 | 0.229 | 0.540 | 206.43 |
| Spat2018 + For + TEve + Herb + (1\|Site) | 222.87 | 0.25 | 0.203 | 0.576 | 202.98 |
| Spat2018 + For + TOP + Herb + SessInv + (1\|Site) | 225.30 | 2.68 | 0.060 | 0.604 | 201.30 |
| Spat2018 + For + TEve + TDiv + Herb + (1\|Site) | 225.63 | 3.01 | 0.051 | 0.600 | 201.63 |
| Spat2018 + For + Herb + SessInv + (1\|Site) | 225.81 | 3.19 | 0.047 | 0.547 | 205.91 |
| (1\|Site) | 226.41 | 3.79 | 0.035 | 0 | 219.36 |
| Spat2018 + (1\|Site) | 227.80 | 5.36 | 0.017 | 0.109 | 217.98 |

For both settlement and recruitment models, fish assemblage variables representing potential foraging impact were stronger predictors of success than observed foraging rates. Herbivore abundance had a strong positive effect on both coral settlement and recruitment, but this effect was greater for recruits (6.62 ± 1.39 SE; Table 3; Figure 4f). Conversely, there was no evidence from either model supporting coral spat or recruit relationships with foraging rate (Table 3; Figure 4). TOP and trait divergence were the strongest predictors of coral settlement success with large negative effects (Table 3; Figure 4c–d). However, the data appears to better support a strong relationship with trait divergence rather than with TOP (Figure 4c). The modelled relationship between coral recruitment and TOP similarly did not appear well-supported by our data, even though this was the largest effect compared with other predictors of recruitment (−7.52 ± 0.30 SE; Table 3; Figure 4h).

**Table 3.**
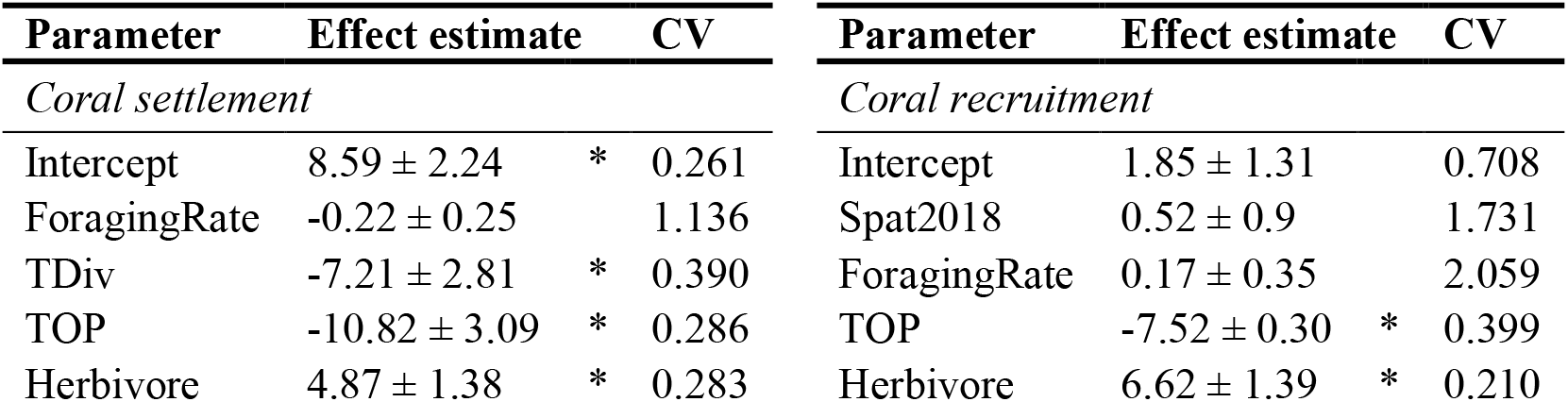
Parameter estimates of selected models exploring the relationship of coral settlement and recruitment with fish assemblage foraging rates, trait divergence (TDiv), trait richness (trait onion peeling index, TOP), and herbivore abundance. Effect estimates are shown with their respective standard error and coefficient of variation. Estimates marked with asterisks (*) are significant *(p* < 0.01).

## Discussion

Our results show that coral settlement and recruitment success are correlated with fish assemblages that have high herbivore abundance but low trait diversity with fewer specialist species present. This aligns with previous studies that suggested the facilitative role of fish assemblages in coral juvenile success and reef recovery through their foraging impacts (Bellwood et al. 2006; Hughes et al. 2007; Cheal et al. 2010; Adam et al. 2011; Rasher et al. 2012). While our results encompass potential foraging impacts and other assemblage indirect effects, our results suggest that fish assemblage diversity could play a role in the conduciveness of a reef environment for coral juvenile growth. We found that herbivore abundance was positively related to coral settlement and recruitment but not as a sole predictor variable. For both recruitment and settlement models, our process of model selection showed that a negative relationship with trait diversity explained variation in coral survival that herbivore abundance could not— trait divergence and richness for settlement and trait richness for recruitment. Of the two trait diversity metrics, divergence best explained the variation in coral settlement patterns (Figure 4; Table 3). There was also little evidence to support relationships with sessile invertivore abundance or foraging rates. While herbivore abundance remains positively associated with coral juvenile survival, we found evidence that this relationship is likely conditional on an assemblage composition that tends to be less trait extreme.

While the modelled relationships with herbivore abundance aligned with our initial prediction, we did not anticipate that its correlation with settlement would be weaker compared with its relationship with coral recruitment. This suggests that coral settlement is more sensitive to differences in trait diversity that is not captured in herbivore abundance. One potential explanation for the differing response to trait diversity in coral settlement to recruitment may be due to recruits having greater energetic stores to overcome or compensate for sub-optimal growth conditions (Ritson-Williams et al. 2009; Doropoulos et al. 2012). This ability to withstand a certain period of sub-lethal inhibition is a likely reason that a wider range of environmental conditions could result in similar recruitment outcomes. Inherent limitations in the temporal matching of our methods may also explain for the differences in fixed effect structures between settlement and recruitment models. Coral settlement and recruitment occur over different temporal scales. The foraging activities most influential for settlement would take place on the scale of weeks before and after summer mass spawning, whereas this would take place on the scale of months to years for recruitment. It is plausible that our sampling duration was more temporally precise for assessing effects on settlement and less aligned for recruitment.

We detected overall stronger effects from assemblage diversity predictors, which represent potential foraging impact, rather than observed foraging rates (Table 3). From both a theoretical and behavioural standpoint, co-occurrence does not necessitate biotic interaction, and so we could not assume all present fish observed were actively foraging in the area (Blanchet et al. 2020). As such, we expected foraging rates to have greater effect sizes than assemblage diversity metrics. The lack of relationship between observed foraging rates and coral settlement and recruitment may be due to highly clustered distributions of foraging sessions, selective patchy foraging across space, or the influence of gregarious foraging behaviours (Hoey and Bellwood 2009; Michael et al. 2013; Streit et al. 2019), resulting in a poor representation of the foraging occurring across each study site.

We recognise that our analyses here are correlative and likely also capture indirect processes that affect coral survival in early life stages in addition to fish assemblage responses to benthic dynamics. The negative relationship between corals and fish trait diversity could point to opposing responses to an external factor we did not examine here such as structural complexity or existing benthic cover. Fish assemblage diversity has been found to be consistently higher when reefs are more structurally complex with increased coral cover (Komyakova et al. 2013; Darling et al. 2017; Richardson et al. 2017; Pombo-Ayora et al. 2020). We focus here on the top-down role of fishes in contributing to conducive environments for corals during settlement and recruitment, but we cannot ignore that the benthic habitat also in turn influences fishes and their foraging behaviour (Vergés et al. 2011; Richardson et al. 2020). It is also possible we detected low settlement and recruitment at sites with increased space pre-emption competition from existing high benthic coverage, which also fostered a more diverse fish assemblage.

Settlement success in this study was associated with fish assemblages that had lower trait divergence (Figure 4b); that is, fewer specialists, even when herbivore abundances were accounted for. This result was in contrast with our hypothesis, and somewhat counterintuitive, because many detritivores located in the centre of our trait space are considered reducers of algal turf sediment load rather than effective substrate-clearing foragers (Purcell and Bellwood 1993; Tebbett et al. 2017). One possible reason for the sensitivity to trait differences in settlement is that trait specialist herbivores may have an initial harmful effect on spat. This negative relationship with trait diversity suggests that the presence of some specialists may have negatively affected survival, whether this was through direct consumption or an indication of other deleterious factors. Spat survival can be negatively correlated to the biomass of grazing fishes (especially parrotfishes) or their feeding scars (Mumby 2009; Baria et al. 2010; Penin et al. 2011b; Trapon et al. 2013a, 2013b).

Excavating and scraping parrotfishes, two feeding modes that are located in the outer extremes of the trait space (Figure 2), have been suggested to be the most disruptive to coral settlement success due to incidental grazing of recently settled corals (Mumby 2009; Trapon et al. 2013b). These grazing fish are often cited as a reason for increased spat survival in small crevices (Nozawa 2012; Brandl et al. 2014; Doropoulos et al. 2016; Gallagher and Doropoulos 2017). Conversely, Brandl et al. (2014) reported positive coral-foraging associations from *Siganus* spp., a group of crevice-feeding algal croppers that are also trait specialists in our study. While our methods were not designed to ascertain relationships from certain species or groups, we do note that algal croppers were abundant at the site with the highest spat counts (North Reef; Figure 2). Despite the risks of incidental grazing mortality, studies find that herbivore abundance and foraging impacts remain beneficial to coral juveniles (Bozec et al. 2015; Graham et al. 2015).

While fewer excavators or scrapers is a likely explanation for increased settlement, we do acknowledge that our study question does not factor how fish foraging impacts on corals may vary in different topographical surroundings. Our use of experimental substrates here likely overestimates the effect of fish-mediated foraging impacts. Because we investigated the relationship between fish and coral assemblages in isolation, we caution against predictive interpretations of the site-wise differences we detected in spat survival with different fish assemblage compositions present. The role of structure in the settlement and recruitment patterns of corals cannot be ignored. Further studies are required to understand how structural complexity mediates this relationship between fish trait diversity and coral settlement.

In this paper, we examine the relationship between fish assemblage diversity and early life stage survival in corals. A conducive habitat is key to coral juvenile survival, and fish could be a part of this environment. While we show here again that herbivore abundance is positively correlated with coral settlement and recruitment success, we highlight that both trait diversity and identity may be important in shaping herbivore effects on coral recruitment. Especially for coral settlement, herbivore abundance is a more “broad stroke” metric compared to trait divergence, which captures potential diminishing returns from specialist foragers. The relationships we found between coral settlement and recruitment and fish trait diversity are one piece of the puzzle that leads to spatial heterogeneity of coral recovery.

## Acknowledgments

We thank the Lizard Island Research Station staff for their support. This study was conducted under a GBRMPA research permit G15/38127.1 valid from 4 December 2015 to 30 January 2022 with ethics approval from the University of St Andrews School of Biology Ethics Committee for non-ASPA research. Funding was provided by the Warman Foundation (to MD and JSM), the John Templeton Foundation (MD, JSM grant #60501 ‘Putting the Extended Evolutionary Synthesis to the Test’), a Royal Society research grant and a Leverhulme fellowship, the Leverhulme Trust Research Centre–the Leverhulme Centre for Anthropocene Biodiversity and a Leverhulme Research Grant (RPG-2019-402, MD), a National Science Foundation–Natural Environment Research Council Biological Oceanography grant (1948946) (JSM, MD), two Ian Potter Doctoral Fellowships at Lizard Island Research Station (DTP and VB) and MASTS small grant to VB. We would also like to thank the anonymous reviewers for their careful feedback on this study.

## Data availability

The associated research data and analysis code can be found in GitHub (github.com/cherfychow/FishTraitCoralRec) with a stable release in Zenodo (doi.org/10.5281/zenodo.7611835) (Chow et al. 2023).

