## Supplementary material for "Coral settlement and recruitment are negatively related to reef fish trait diversity"

in *Coral Reefs*

**Authors:**

Cher F Y Chow, Caitlin Bolton, Nader Boutros, Viviana Brambilla, Luisa Fontoura, Andrew S Hoey, Joshua S Madin, Oscar Pizarro, Damaris Torres-Pulliza, Rachael M Woods, Kyle J A Zawada, Miguel Barbosa, Maria Dornelas

**Corresponding author:**

Cher Chow

Centre for Biological Diversity, School of Biology, University of St Andrews

### Assigning foraging mode categories

Functional group classifications roughly summarise the feeding method, substrate impact, jaw morphology, and dentition traits. Six of these are exclusive only to the herbivore trophic group: (1) excavators, which are parrotfishes (Labridae, Scarini) that remove substrate with bites; (2) scrapers consist of *Scarus* spp. and *Hipposcarus* spp. parrotfishes that scrape epilithic algal turf from substrate but do not excavate substrate with bites; (3) croppers remove turf algae biomass but leave the holdfasts at the base intact; (4) browsers similarly leave holdfasts intact but remove macroalgae tissue; (5) brushers are grazing detritivores that have brush-like teeth to remove particles around turf algae but do not remove algal tissue; and (6) farmers, which are highly territorial damselfishes (Pomacentridae) that selectively graze and cultivate algae stands within their territories. Pickers, suction feeders, ambush feeders, and active feeders are not exclusive to trophic groups. Pickers also feed benthically but for fish that selectively remove their prey from benthic cover while foraging. Suction feeders refers to planktivores and some piscivores, like the sling-jawed wrasse, *Epibulus insidiator*. Ambush and active feeders are distinguished by the way they approach prey, where active feeders employ pursuit and ambush feeders do so by sudden bursts.

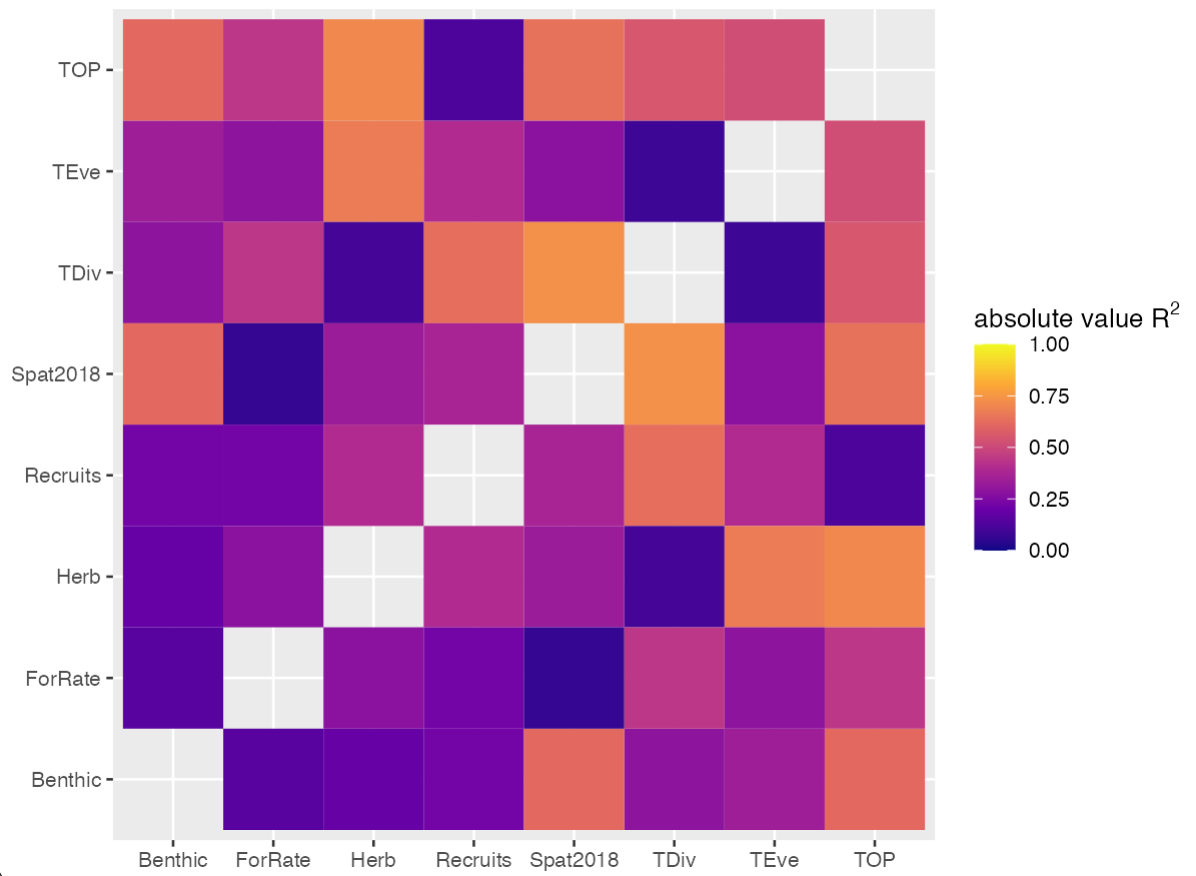
Because information within FishBase varied across species, we allocated values from other literature sources and RUVs for species with incomplete data (**Table 2.1**). *Thalassoma nigrofasciatum* and *Chlorurus spilurus* were data deficient species due to relatively recent taxonomic reviews. For these two species, I used data associated with their previous synonym or closest relative (*Thalassoma jansenii* and *Chlorurus sordidus* respectively) and validated these trait values with observed behaviour in RUVs.

**Figure S1**. Collinearity check for coral settlement and recruitment model predictor variables. The pairwise Pearson correlation coefficients are shown represented shaded by a colour scale. Absolute values closer to *r =* 1.00 are shown with lighter shading. We accepted predictor variables if they were below a threshold of *r* = 0.80, where the maximum here is *r* = -0.73 between trait divergence (TDiv) and 2018-19 spat counts (Spat2018).

### Trait space construction

We used scree plot visualisations of resulting eigenvalues to determine an optimum number of dimensions to preserve for functional diversity calculations (**Figure S2a**). The resulting global trait space consisted of four axes with a trait space quality, or proportion of sum of eigenvalues, of 0.366 (Villéger et al. 2008). We also validated the trait space quality by comparing the reduced space dissimilarities with original dissimilarities in a Mantel test (*r* = 0.868, *p* < 0.01) and a Shephard diagram (**Figure S2b**).

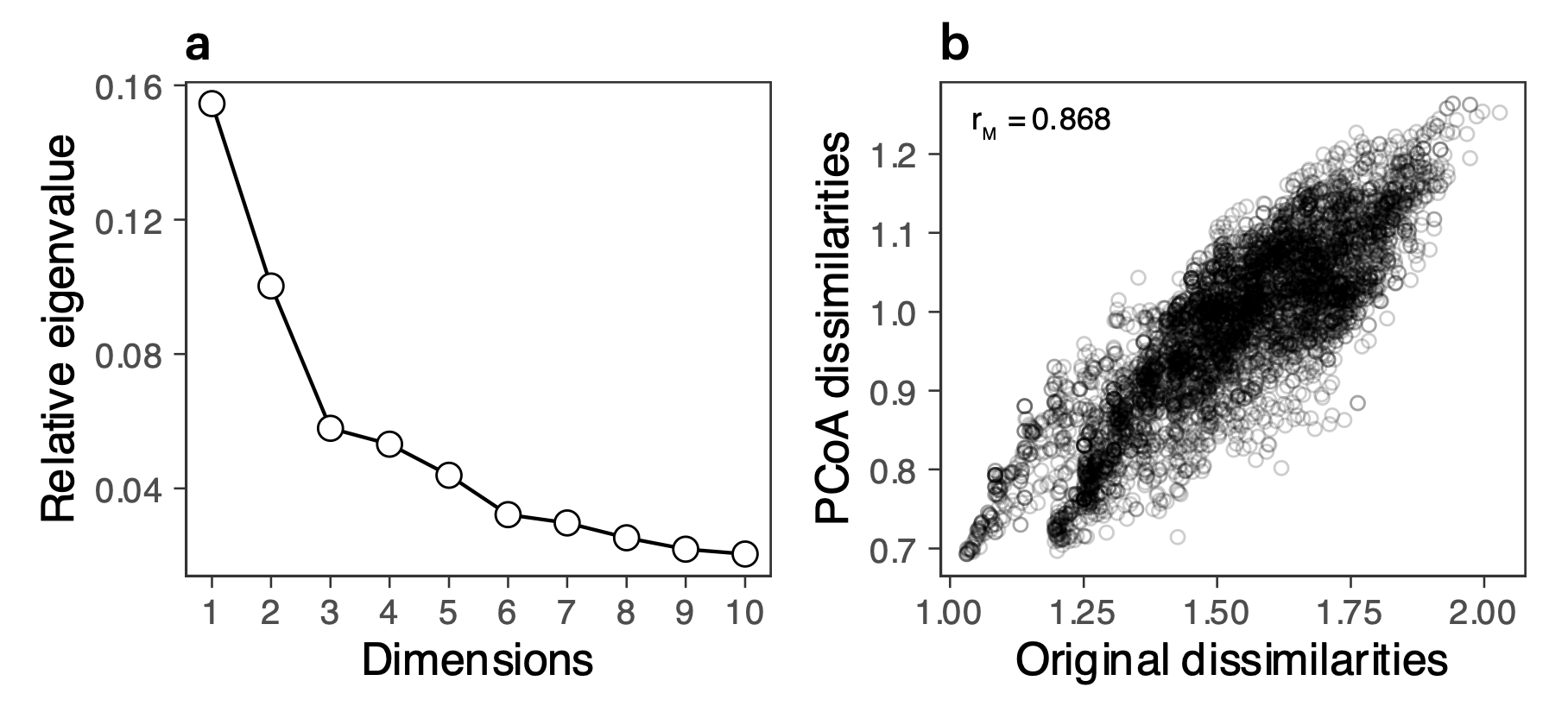

**Figure S2.** Principal coordinates analysis (PCoA) diagnostics. A scree plot (a) shows the relative eigenvalues, representing proportion of variation explained, for the first 10 principal coordinates with the characteristic “elbow” that indicates optimum coordinates to preserve in the trait space. A Mantel test (b) validates the PCoA results as well by correlating original dissimilarities with the scaled PCoA dissimilarities to assess how the dimension reduction preserves the variation represented in the original data (r_M_ = 0.868, *p* < 0.01).

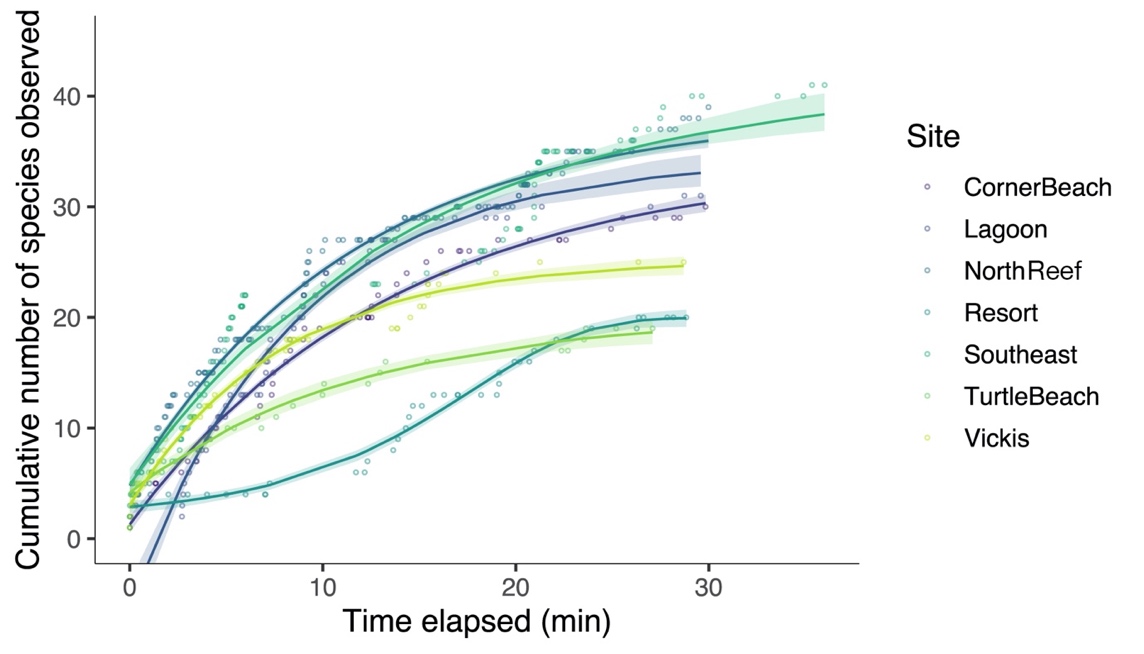

**Figure S3.** Species accumulation over a 30-minute sampling duration in remote underwater video observations of fish assemblages conducted at seven study sites. Time elapsed for Southeast appears to exceed 30 minutes due to an omitted 6-minute period caused by boat disturbance. Each point represents a timestamp at which fish individuals were observed. An asymptotic or Gompertz nonlinear regression model was fitted for each site to determine the species accumulation curve, illustrated here as lines with bootstrapped 95% confidence intervals shown in shaded ribbons. All sites fitted an asymptotic model except for Resort where a Gompertz equation fitted best.

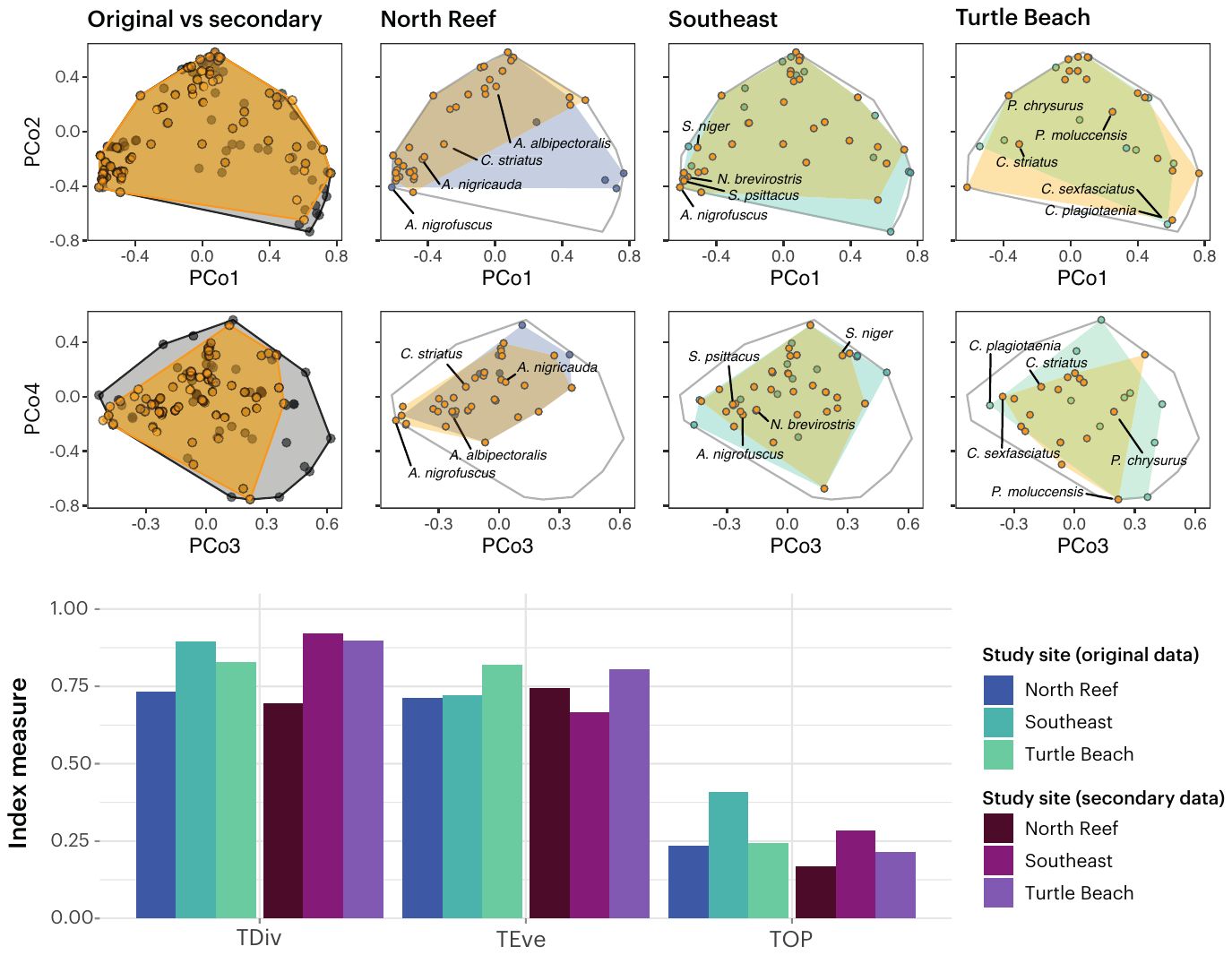

**Figure S4**. Trait space sensitivity analysis of changes in fish assemblages from additional video fields of view. At the top, comparisons of the trait spaces are shown where each circle represents a species. Fish assemblage trait spaces observed in additional second videos are shaded in orange, while original fish assemblage trait spaces used in this study are coloured by study site (blues). The first panel column is a pooled aggregate of the assemblages from all three study sites used in this sensitivity analysis, comparing the total differences between the original video observations with secondary videos (orange). The three highest ranking species in terms of abundance are indicated with text labels. The pooled fish assemblage for all sites from all videos is shown outlined in light grey. Comparisons of the trait diversity metrics which describe these trait spaces are shown at the bottom where sites from the secondary data are shown in shades of plum.

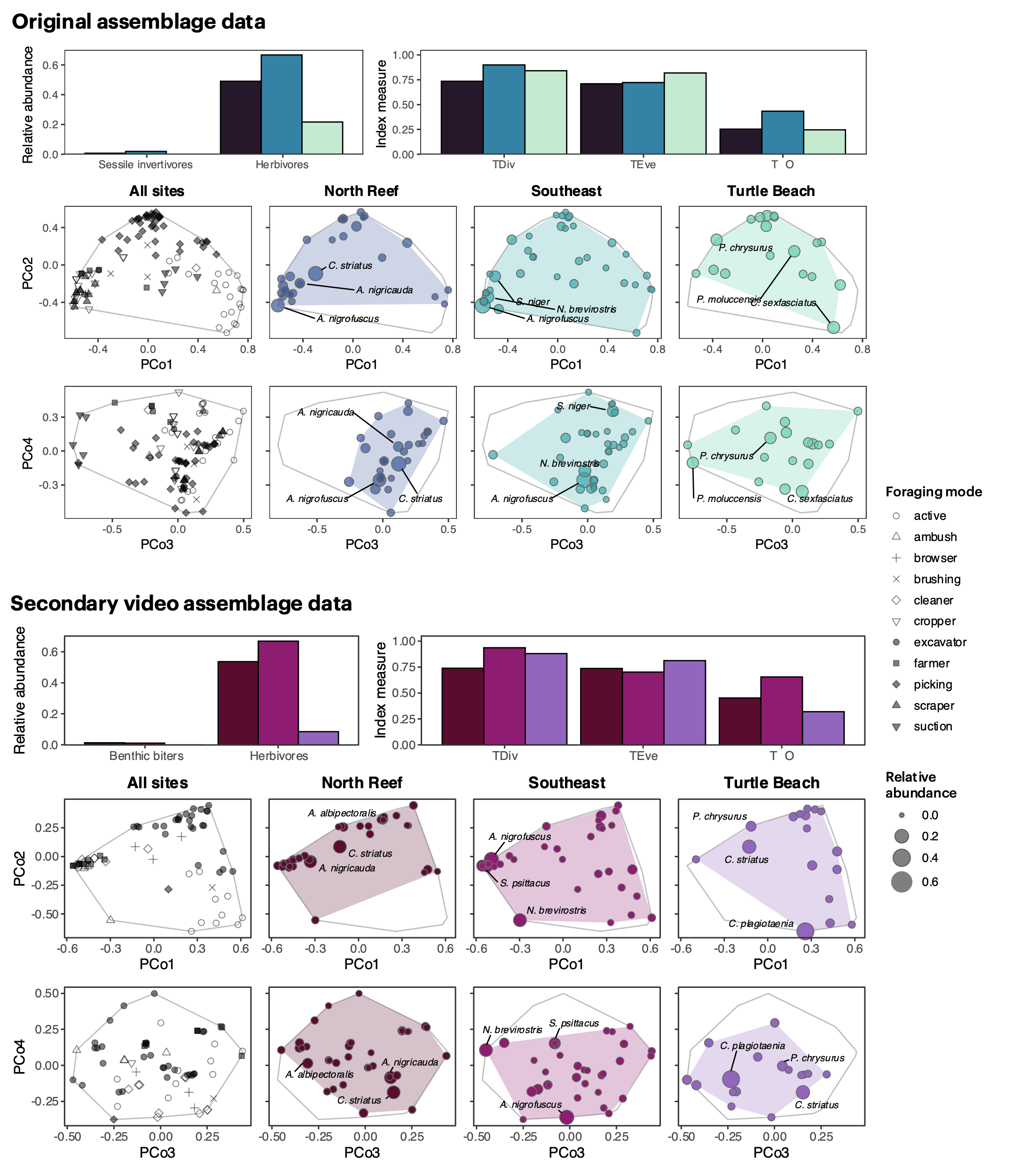

**Figure S5.** Independent comparison of the trait diversity of fish assemblages observed in two sets of remote underwater videos. The top half of this figure represents the original observations from the primary videos, and the bottom half represents observations from the secondary set of videos used for the sensitivity analysis. These trait spaces were constructed independently of each other to determine whether the scaling would reflect a similar ordination pattern and trait diversity indices. The bar graphs show measures for relative benthic biter and herbivore abundance (top left) and trait diversity indices (top right): trait divergence (TDiv), trait evenness (TEve), and trait onion peeling index (TOP). The arrays shown are a four-dimensional representation of assemblages according to six foraging traits of species. Species are represented by circles, with varying sizes by relative abundance. Distance between circles represents trait dissimilarity between species.

### Trait-based foraging rates

We derived the trait-weighted coefficient (S*_i_*) for species foraging rates from a modified version of the trait table. Trophic group, foraging mode, and water column position were expressed as ordered factors to reflect bite impact. Within each trait, we ordered factors according to ability to remove coral-inhibiting biogenic cover as supported by previous studies on trophic differences in bites (Green and Bellwood 2009). We then computed Euclidean distances between all species according to these modified traits and performed a Principal Components Analysis (PCA). The resulting first principal component (PC1) explained 0.898 of the variation in the distances. We validated this principal component by ensuring that species rankings reflected the factor ordering in the trait table. If it was scaled in the inverse, we corrected this ranking by multiplying the PC1 value by -1. **Equation S1** shows the scaling of this principal component by its range to derive the trait-weighting factor.

**Equation S1**

$$S_{i}=0.01 \frac{\left( {PC1}_{i}+ {PC1}_{\text{min}}+0.1 \right)}{2}$$

**Table S1**. Trait-weighted coefficients used in the scaling of bite rates to foraging rates that estimate foraging impact for biting species. These factors are generated according to three functional traits of trophic group, foraging mode, and water column position of feeding. Larger factors represent species that deliver greater feeding impacts on reef benthos, and vice versa for lower factors.

| **Species** | **TWF** | **Species** | **TWF** |
| --- | --- | --- | --- |
| *Chlorurus microrhinos* | 3.67 | *Halichoeres melanurus* | 1.58 |
| *Chlorurus spilurus* | 3.67 | *Hemigymnus melapterus* | 1.58 |
| *Naso lituratus* | 3.33 | *Thalassoma hardwicke* | 1.58 |
| *Hipposcarus longiceps* | 3.33 | *Thalassoma lunare* | 1.58 |
| *Scarus altipinnis* | 3.33 | *Chaetodon baronessa* | 1.56 |
| *Scarus frenatus* | 3.33 | *Chaetodon lunulatus* | 1.56 |
| *Scarus niger* | 3.33 | *Chaetodon melannotus* | 1.56 |
| *Scarus rubroviolaceus* | 3.33 | *Chaetodon auriga* | 1.22 |
| *Acanthurus nigrofuscus* | 2.98 | *Chaetodon citrinellus* | 1.22 |
| *Siganus corallinus* | 2.98 | *Chaetodon vagabundus* | 1.22 |
| *Siganus puellus* | 2.98 | *Pomacanthus sexstriatus* | 1.22 |
| *Zebrasoma scopas* | 2.98 | *Pomacentrus chrysurus* | 1.22 |
| *Dischistodus prosopotaenia* | 2.98 | *Sufflamen chrysopterum* | 1.22 |
| *Pomacentrus wardi* | 2.98 | *Stethojulis trilineata* | 1.18 |
| *Stegastes apicalis* | 2.98 | *Chrysiptera caesifrons* | 1.17 |
| *Ctenochaetus striatus* | 2.28 | *Pomacentrus moluccensis* | 1.17 |
| *Parupeneus barberinus* | 1.92 | *Acanthochromis polyacanthus* | 0.05 |
| *Halichoeres marginatus* | 1.58 |  |  |

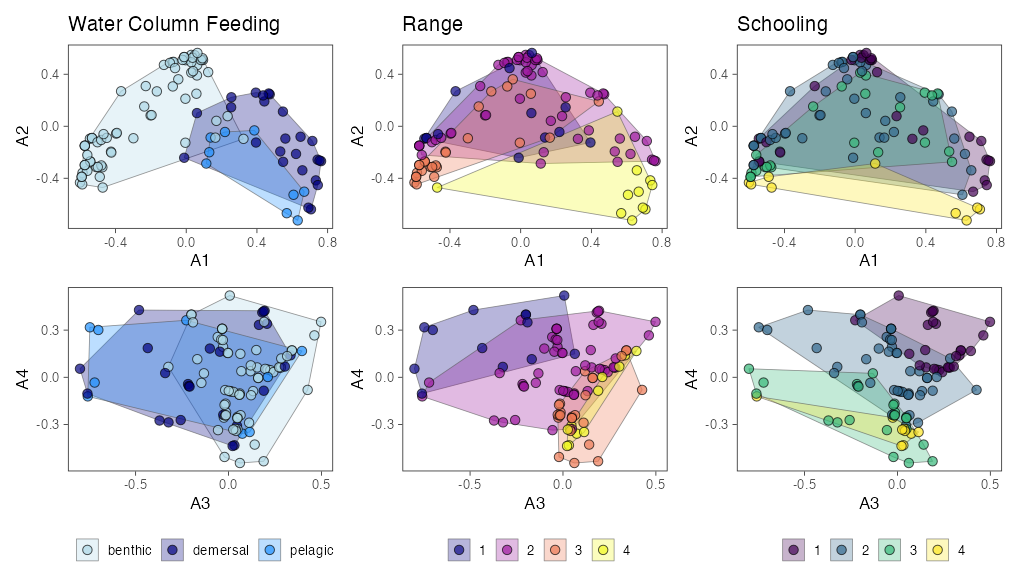

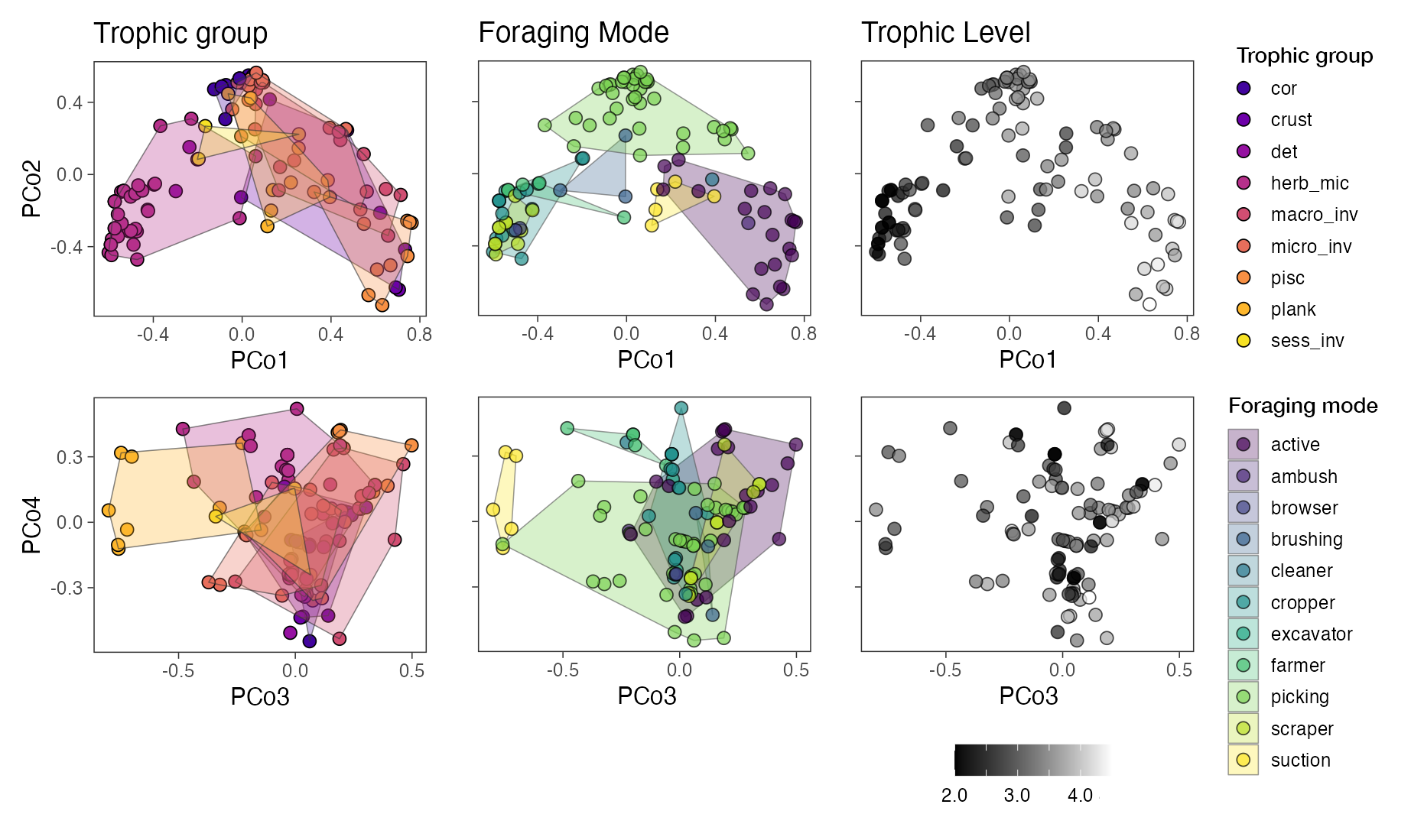
**Figure S6**. Four-dimensional trait space clustering patterns according to three functional traits of trophic group, foraging mode, and trophic level. Each point represents an observed species within this trait space. The distance between points represents relative dissimilarities. Points are grouped according to the levels within each trait, where levels are distinguished by colour and outlined with a convex hull where possible.

**Figure S7**. Four-dimensional trait space clustering patterns according to three functional traits of water column position of feeding, active range size, and schooling. Range size is ordered from 1-4, where 1 represents highly territorial species and 4 pelagic, wide-ranging species. Schooling is ordered from 1-4, where 1 represents solitary species to 4 representing species forming large schools. Each point represents an observed species within this trait space. The distance between points represents relative dissimilarities. Points are grouped according to the levels within each trait, where levels are distinguished by color and outlined with a convex hull.

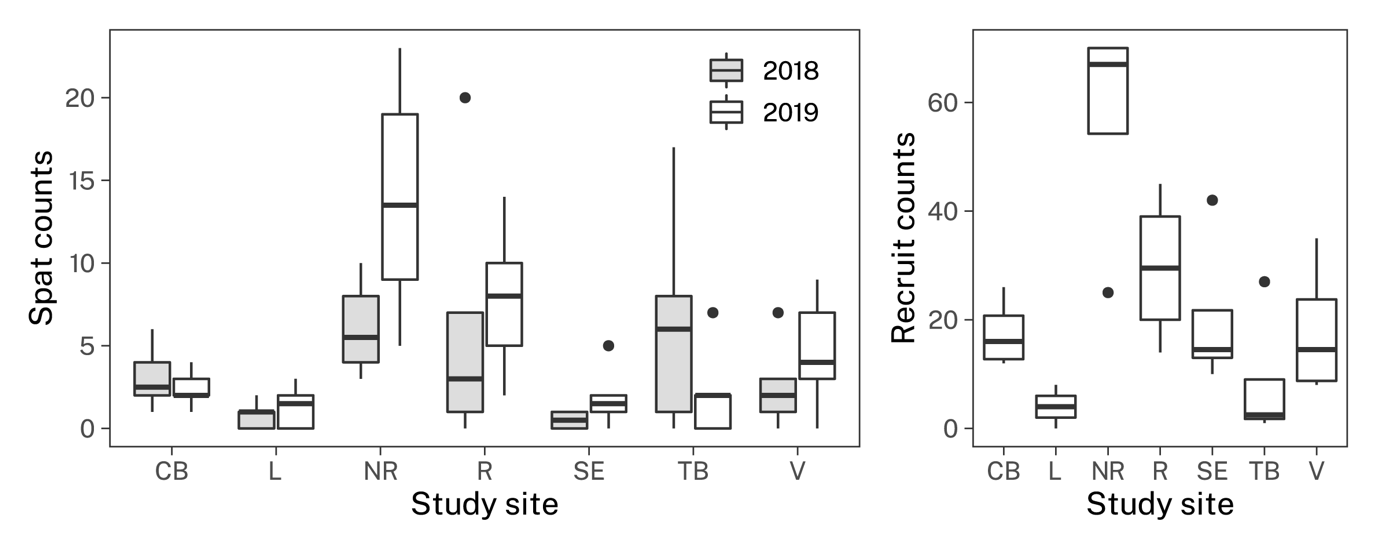
**Figure S8.** Site differences in coral settlement and recruitment according to spat and recruit counts. Recruit counts are for only 2019. Spat counts came from six experimental tiles and recruit counts from four photomosaics of reef sections. Boxes represent the distribution of these counts, where 50% of the distribution is contained within the box and the central line at the mean. Box whiskers span the range of the data with outliers shown as black circles.

**Table S2**. Relative species contributions to trait diversity indices at each study site. These relative species contributions are generated by omitting species sequentially from trait space construction without affecting pairwise dissimilarities. Mathematically it represents the difference in original and species-omitted trait indices divided by the site’s original trait index. We show here the top 2 contributing species by absolute value to each trait diversity index for each study site.

| **Site** | **Species** | **TEve** | **TDiv** | **TOP** |
| --- | --- | --- | --- | --- |
| Corner Beach | *Pomacentrus moluccensis* |  |  | 0.191 |
|  | *Scarus niger* |  |  | 0.061 |
|  | *Hemigymnus melapterus* |  | -0.024 |  |
|  | *Uraspis uraspis* |  | 0.003 |  |
|  | *Siganus doliatus* | -0.029 |  |  |
|  | *Hipposcarus longiceps* | 0.024 |  |  |
| Lagoon | *Plectropomus leopardus* |  |  | 0.157 |
|  | *Acanthochromis polyacanthus* | -0.028 |  | 0.113 |
|  | *Thalassoma lunare* |  | -0.021 |  |
|  | *Plectropomus leopardus* |  | -0.020 |  |
|  | *Hemigymnus melapterus* | -0.036 |  |  |
| North Reef | *Plectropomus leopardus* |  |  | 0.140 |
|  | *Thalassoma nigrofasciatum* |  |  | 0.131 |
|  | *Ctenochaetus striatus* |  | -0.175 |  |
|  | *Acanthurus nigrofuscus* | -0.034 | 0.091 |  |
|  | *Naso brevirostris* | -0.028 |  |  |
| Resort | *Pomacentrus moluccensis* |  |  | 0.224 |
|  | *Lutjanus carponotatus* |  | 0.022 | 0.222 |
|  | *Hemigymnus melapterus* |  | -0.025 |  |
|  | *Siganus punctatissimus* | -0.040 |  |  |
|  | *Siganus punctatus* | -0.040 |  |  |
| Southeast | *Grammatorcynus bicarinatus* |  |  | 0.160 |
|  | *Abudefduf whitleyi* |  |  | 0.144 |
|  | *Acanthurus nigrofuscus* |  | 0.038 |  |
|  | *Naso brevirostris* |  | 0.013 |  |
|  | *Ctenochaetus striatus* | 0.031 |  |  |
|  | *Cheilinus trilobatus* | 0.051 |  |  |
| Turtle Beach | *Caranx sexfasciatus* |  | 0.033 | 0.294 |
|  | *Parupeneus cyclostomus* | -0.022 |  | 0.158 |
|  | *Scolopsis bilineata* |  | 0.034 |  |
|  | *Cheilinus chlorourus* | -0.054 |  |  |
| Vicki’s | *Cephalopholis argus* |  |  | 0.159 |
|  | *Pomacentrus moluccensis* |  |  | 0.088 |
|  | *Pomacentrus grammorhynchus* |  | 0.029 |  |
|  | *Zebrasoma scopas* | -0.040 | 0.039 |  |
|  | *Lutjanus gibbus* | -0.029 |  |  |
